# Nighttime sleep promotes mechanosensory habituation in *Drosophila*

**DOI:** 10.1101/2025.11.17.688779

**Authors:** Simon A. Lowe, Abigail D. Wilson, Ko-Fan Chen, James E.C. Jepson

## Abstract

Sleep is a highly conserved behaviour that facilitates episodic, procedural, and/or associative modes of memory across species ^1-3^, a process hypothesised to involve structural downscaling of synapses potentiated during waking episodes ^4-6^. Habituation, an ancient form of non-associative memory in which repetition of a stimulus leads to diminishing behavioural responses, is considered a prerequisite for such complex forms of learning ^7,8^. Yet whether sleep also influences the capacity of organisms to habituate is poorly understood. Here we examine this question using the fruit fly, *Drosophila*. We describe an automated system that yields long-term analogue measurements of habituation to mechanosensory stimuli in adult flies. Using this platform, we find that *Drosophila* lacking neuronal calcium sensor Neurocalcin – which exhibit reduced sleep specifically during the night ^9^ – display impaired mechanosensory habituation during the day. Mimicking a method used to treat insomnia ^10^, we show that compressing nighttime duration both restores consolidated night sleep and renormalises habituation in *Neurocalcin* mutants. Conversely, in wild type flies without sleep deficits, decreasing night length reduces total night sleep and disrupts daytime habituation. In wild type flies, exposure to longer nights – which increases overall night sleep – enhances synaptic downscaling in sensory circuits. Strikingly, this effect is abolished in *Neurocalcin* mutants, which do not exhibit increased sleep under these conditions. Collectively, our findings reveal a role for nighttime sleep in facilitating habituation and suggest that this may occur by promoting synaptic downscaling.

## RESULTS AND DISCUSSION

### Induction of mechanosensory habituation in adult *Drosophila*

Numerous assays have been developed to induce habituation of adult *Drosophila* to visual, olfactory, and gustatory, stimuli ^11-14^. However, while recently examined in *Drosophila* larvae ^15^, habituation to mechanical stimuli has been comparatively understudied in adult fruit flies. More broadly, whether sleep quality modulates the ability of adult flies (or other organisms) to habituate to environmental stimuli remains an open question.

We therefore sought to investigate potential links between sleep and habituation to mechanosensory stimuli in adult *Drosophila*. The synaptic homeostasis hypothesis stipulates that sleep facilitates downscaling of synapses potentiated during waking experience, thus enabling memory by preventing neuronal hyper-metabolism ^16^ and enhancing signal-to-noise ratios across circuits ^5^. Ongoing locomotor activity occurring during wakefulness could induce an opposite change, potentiating synaptic connections in circuits mediating movement ^17,18^ and mechanosensory feedback ^19-21^. We therefore hypothesised that enhancing this potentiation by limiting sleep (thus prolonging locomotion) might reduce sleep-induced synaptic downscaling and impair the ability of habituation-promoting inhibitory mechanisms ^22,23^ to suppress motor circuit activity in response to repeated mechanical stimuli.

To test this hypothesis, we developed a simple, high-throughput habituation assay using the DART (*Drosophila* ARousal Tracking) system (Fig. 1A) ^24^, which combines continuous video-tracking of individual flies with mechanical stimuli delivered via platform-associated motors. DART can be programmed to deliver consistent mechanosensory stimuli that elicit robust, transient, locomotor responses in control flies (Figure S1A), the magnitude and dynamics of which can be quantified. We repurposed the DART system to test for mechanosensory habituation in adult flies by delivering multiple identical stimuli at intervals of 10 min (Fig. 1B). Flies responded to repeated mechanical stimuli with locomotor responses that gradually attenuated in magnitude (Fig. 1C), quantified as the mean peak speed in the 1 min following each stimulus (Fig. 1D).

**Fig. 1.**
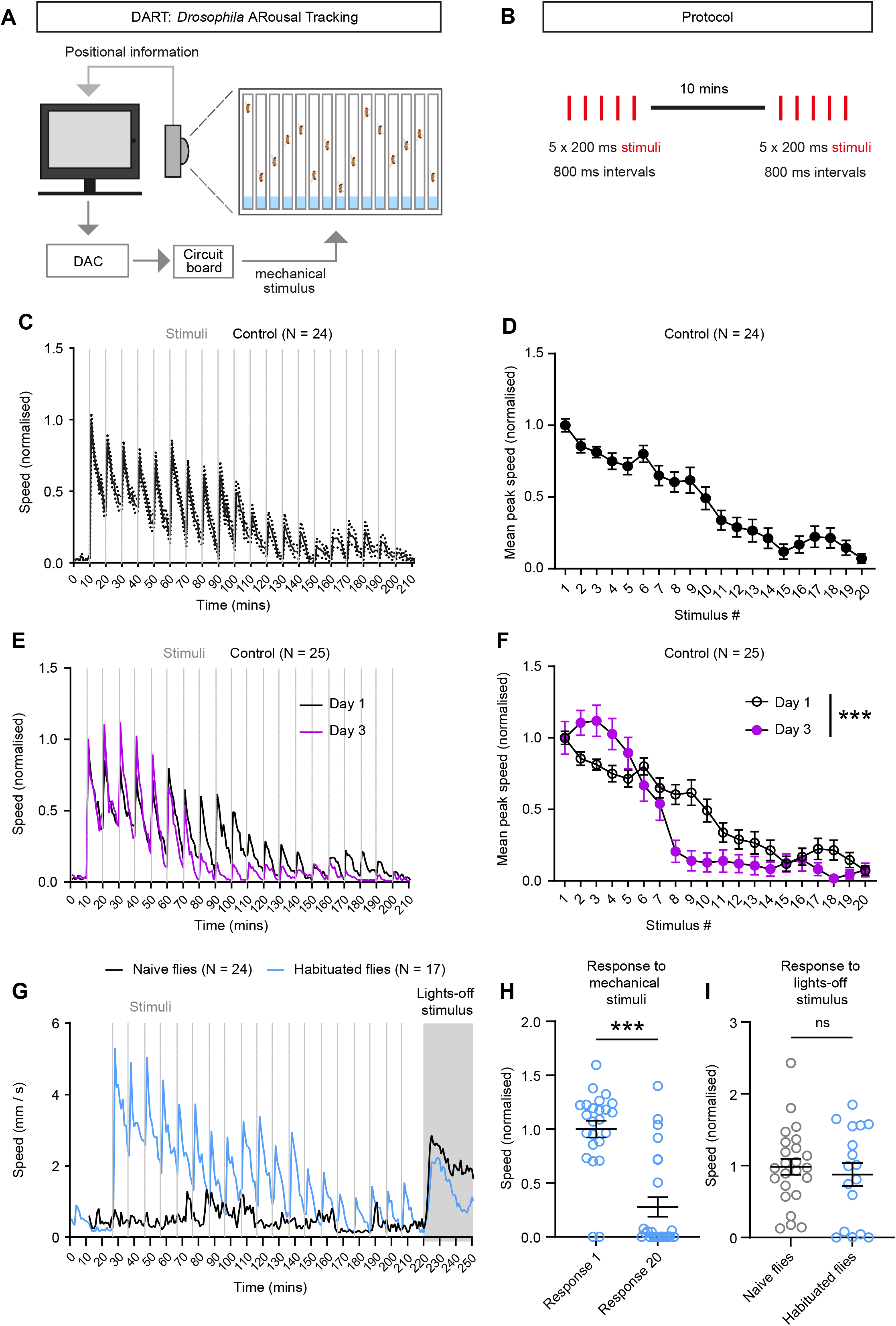
Mechanosensory habituation in adult *Drosophila*. (**A**) Schematic illustrating the *Drosophila* ARousal Tracking (DART) system. (**B**) Protocol used to induce mechanosensory habituation. (**C**) Mean speed over time in one-minute bins, and (**D**) mean speed in the bin following each stimulus, of N = 24 control (iso31) male flies, exposed to 20 successive mechanical stimuli. Data are normalised to the mean in the bin following the first stimulus, i.e. the naïve response. (**E-F**) Mean speed of locomotion over time (E) and mean response in the one-minute bin following each stimulus (F) in N = 25 control adult male flies, in response to an identical habituation protocol applied on three consecutive days. Data from Day 1 and Day 3 are shown. Data are normalised to the response to the first stimulus within each day. Interaction between stimulus number and day: ***p<0.0005, two-way ANOVA. (**G**) Mean speed of locomotion over time as a “lights-off” stimulus is delivered to naïve flies (black, N = 24) and immediately following a train of 20 mechanosensory stimuli (blue, N = 17). (**H**) Normalised locomotor responses of control adult male flies to the 1^st^ and 20^th^ mechanical stimulus shows a clear decline in response to repeated stimulus: N = 17. ***p<0.0005, Mann-Whitney U-test. (**I**) Response to lights-off does not differ between naïve flies (N = 24) and those habituated to the distinct mechanosensory stimuli (N = 17). p = 0.7187, Mann-Whitney U-test. Error bars: standard error of the mean (SEM).

If this decrement represents true habituation rather than generalised sensory adaptation, fatigue, or structural damage, it should exhibit key properties that define habituation: spontaneous recovery, potentiation of habituation over repeated bouts, and stimulus-specificity ^7,8^. We sought to validate the assay by testing for these characteristics.

After attenuation of locomotion following 20 mechanical stimuli, locomotor speeds exhibited spontaneous recovery over time in response to single mechanical stimuli, with ∼ 80% recovery occurring following 24 h (Figure S1B-F). We found that the rate of mechanosensory habituation could be significantly enhanced by applying a habituation-inducing paradigm of 20 identical mechanical stimuli on three consecutive days (Fig. 1E,F). Finally, after 20 consecutive mechanical stimuli, we delivered a distinct ‘lights-off’ visual stimulus expected to similarly induce a startle response (Fig. 1G). Despite the robust decrease in locomotor response to repeated mechanical stimuli (Fig. 1G,H), locomotor responses to the visual stimulus did not differ in magnitude between naïve and mechanically habituated flies (Fig. 1G,I). Hence, progressive reductions in locomotor responses in our paradigm are consistent with well-defined characteristics of habituation ^7,8^.

A number of platforms analogous to DART can track locomotor activity of populations of flies using video- or infrared beam-based analysis, and can deliver mechanosensory stimuli or be modified to do so. These include the *Drosophila* Activity Monitor ^25^, Ethoscope, ^26^ and Zantiks ^27^ systems. We designed a similar habituation protocol using the Zantiks automated-tracking system (Figure S1F; see Methods). The Zantiks system could detect and quantify robust locomotor responses of adult flies to single motor-driven stimuli (Figure S1G), as well as a progressive decline in locomotor response to repeated, identical mechanosensory stimuli that was qualitatively similar to that observed using the DART system (Figure S1H, I and Fig. 1D). These data provide cross-platform validation of our findings using the DART.

### Neurocalcin promotes mechanosensory habituation in *Drosophila*

Similarly to humans, deep sleep in *Drosophila* primarily occurs during the night ^28^. We therefore sought to test whether night sleep influences mechanosensory habituation induced during the day. Mechanically disrupting sleep provides a potential means to do so. However, homeostatic rebound sleep induced by sleep deprivation ^29^ could potentially cause confounding changes in stimulus-induced movement during the day. Hence, we utilised a genetic manipulation that selectively disrupts night sleep whilst leaving daytime sleep intact: loss of the highly conserved neuronal calcium sensor Neurocalcin ^9^.

In contrast to mutations such as *shaker, sleepless, fumin*, and *insomniac* – which disrupt both day and night sleep in *Drosophila* ^30-34^ – loss of Neurocalcin specifically impairs sleep during the night in oscillating light: dark conditions ^9^, providing a means to examine the relationship between night sleep and daytime habituation. Indeed, using our paradigm to measure the rate and amplitude of mechanosensory habituation in control flies and flies homozygous for a *Neurocalcin* null allele (*Nca*^KO1^) ^9^, we found that loss of Neurocalcin significantly reduced the degree of mechanosensory habituation, an effect that was particularly prominent after ∼ 10 mechanical stimuli (Fig. 2A,B). We observed similarly reduced mechanosensory habituation in flies homozygous for an independent *Neurocalcin* null allele (*Nca*^KO2^) (Figure S2A-B), confirming the above genotype-phenotype correlation.

**Fig. 2.**
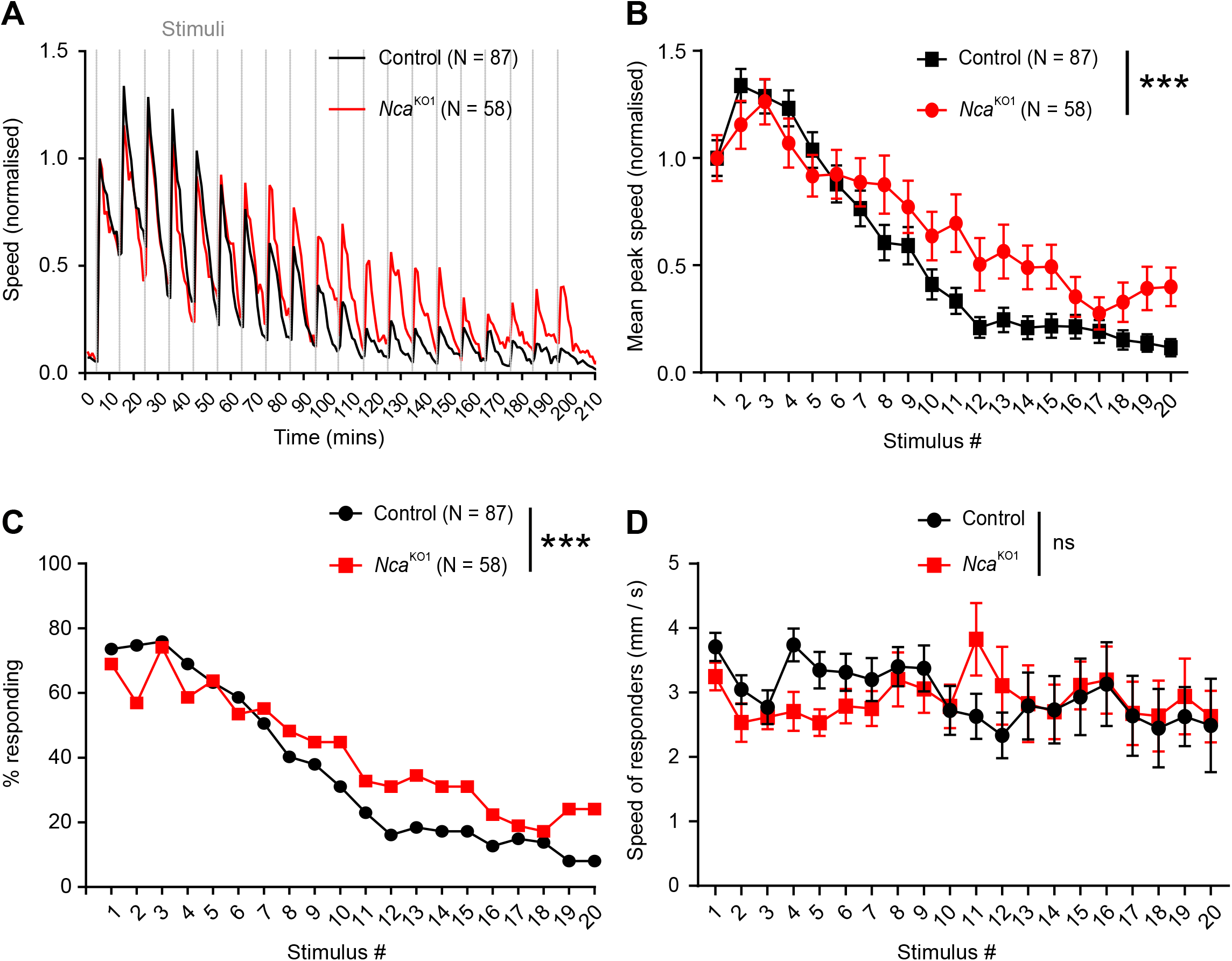
*Neurocalcin* knockout flies exhibit impaired mechanosensory habituation. (**A**) Mean speed of locomotion over time, and (**B**) mean speed in one-minute bin following each stimulus of control (N = 87) and *Nca* knockout (*Nca*^KO1^) (N = 58) adult male flies exposed to 20 consecutive mechanosensory stimuli. Data are normalised within each genotype to the naïve response. *Nca* knockout flies habituate more slowly than controls (interaction between stimulus number and genotype: p<0.0001, two-way ANOVA). Error bars: SEM. (**C**) Proportion of flies “responding” decreased over multiple stimuli, with the rate of decrease slower for *Nca*^KO1^ flies. (**B**) Mean speed of flies “responding” did not decrease in response to repeated stimuli, nor did it differ between genotype (main effect of stimulus number: p=0.2407, and of genotype: p=0.3761; interaction between stimulus and genotype: p=0.4639, two-way ANOVA). Error bars: SEM.

Neurocalcin could potentially modulate mechanosensory habituation by enhancing sensitivity to mechanical stimuli. We therefore tested locomotor responses in control and *Neurocalcin* knockout flies to mechanical stimulus strengths ranging from 1% to 100% of the strength used in the habituation protocol. *Neurocalcin* knockout flies showed statistically similar amplitudes of locomotor responses to isogenic controls at 6 out of 7 stimulus strengths tested, and a slightly enhanced response to stimuli 10% of the strength used in the habituation protocol (Figure S2C). These data therefore did not support a general increase in mechanosensory sensitivity in *Neurocalcin* knockout flies, and confirmed that the habituation protocol utilised a stimulus strength that was beyond the plateau at which there was no difference between control and *Neurocalcin* knockout flies.

Since our habituation protocol yielded long-term quantitative recordings of individual flies, we were able to examine whether the reduction in habituation in *Neurocalcin* knockout flies was due to a change in number of flies responding to mechanical stimuli, or a decrease in the magnitude of the individual flies’ responses. To separate response to a stimulus from background noise, we defined a threshold for a “response” to each stimulus as ≥10% of the response to the mean naïve response. We found that for both control and *Neurocalcin* knockout flies, the attenuation in response came entirely from the proportion of flies responding (Fig. 2C). When only “responding” flies were included in analysis, there was no decrease in the amplitude of locomotor responses across repetitive stimuli, nor any difference between genotype (p = 0.46, main effect of Genotype & of Stimulus #, 2-way ANOVA) (Fig. 2D). The all-or-nothing nature of locomotor responses in our paradigm supports the premise that the decrease in mean amplitude across repeated stimuli represents a learning event i.e an altered “decision” of whether to respond or not to a given stimulus, as opposed to a motor defect affecting the ability to sustain robust reactions over multiple stimuli.

### Short nights restore consolidated sleep and mechanosensory habituation in *Neurocalcin* knockout flies

Hippocalcin, the mammalian homologue of Neurocalcin, promotes synaptic plasticity and memory in mice ^35-39^, and cognitive development in humans ^40,41^. Thus, we sought to test whether Neurocalcin facilitates habituation through its effect on night sleep, or via sleep-independent processes potentially linked to memory.

We previously documented that night sleep loss in *Neurocalcin* mutant flies was particularly prominent under long night, winter-like, conditions ^9^, suggesting that night length can modulate sleep loss in *Neurocalcin* mutant flies. Interestingly, increasing sleep drive by restricting the period of consolidated night sleep is utilised in cognitive behavioural therapy for insomnia, and has similarly been shown to rescue sleep loss in a variety of *Drosophila* sleep mutants ^10^. We therefore examined sleep architecture in control and *Neurocalcin* knockout flies under short and long night conditions using a multibeam monitoring system that facilitates high-resolution quantification of sleep in *Drosophila* ^42^ (Figure S3A).

Control and *Neurocalcin* knockout flies were raised under standard 12 h light: 12 h dark conditions then, after 3-5 days of adulthood, were shifted to three days of long (8 h light: 16 h dark) or short (16 h light: 8 h dark) night conditions, with sleep quantified on the third day of each light: dark cycle. Consistent with prior results ^9^, we confirmed reduced nighttime but not daytime sleep in *Neurocalcin* knockout flies under long night conditions (Fig. 3A,C and Figure S3B-C), and found that this decrease was associated with fragmentated sleep, characterised by a significant increase in the number of sleep bouts and a concomitant reduction in sleep bout duration (Fig. 3D,E). In contrast, under short night conditions, overall night sleep (Fig. 3B,C) and the degree of sleep consolidation (Fig. 3D,E) were not significantly different between *Neurocalcin* knockout flies and controls. Day sleep was also slightly increased in *Neurocalcin* knockout flies under short night conditions, likely due to reduced locomotor activity towards lights-off (Fig. 3B,Figure S3B-C). Thus, changing night length provides a means to tune night sleep quality on-demand in *Neurocalcin* knockout flies, allowing us to disentangle sleep-dependent versus -independent functions of Neurocalcin.

**Fig. 3.**
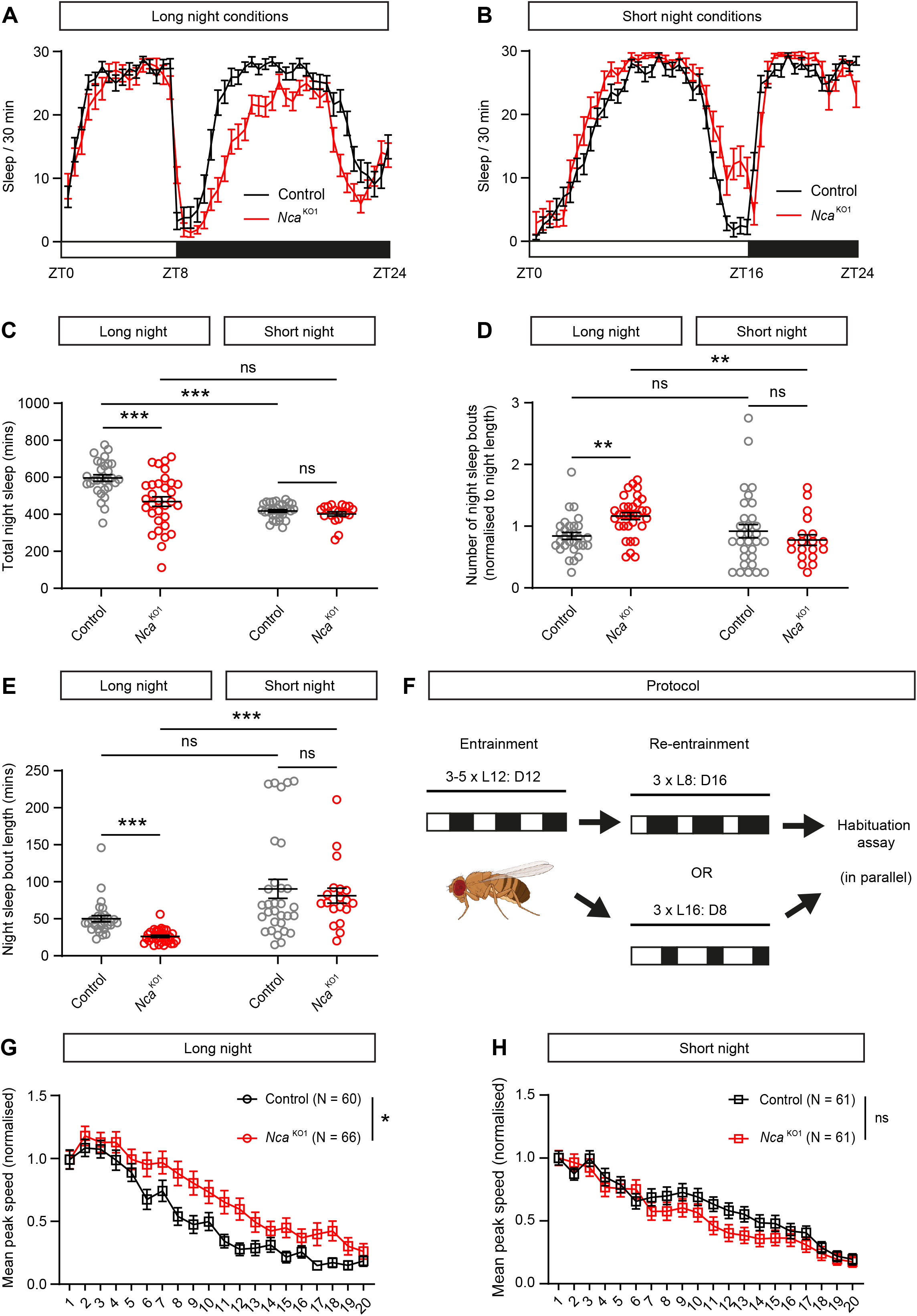
*Neurocalcin* modulates mechanosensory habituation through a sleep-dependent mechanism. (**A-B**) Mean sleep levels of adult male flies over 24 h under (A) short day, long night (L8: D16) conditions (control: N = 32; *Nca*^KO1^: N = 32), and (B) long day, short night (L16: D8) conditions (control: N = 31; *Nca*^KO1^: N = 20). (**C-E**) Total minutes of sleep during subjective night (C), number of individual sleep bouts occurring during the night normalised to hours of night (D), and mean length of sleep bouts occurring during the night (E). Kruskal Wallis test with Dunn’s multiple comparison test. (**F**) Protocol for exposure to short or long night conditions prior to habituation assay. (**G-H**) *Nca*^KO1^ flies habituate more slowly than controls in (G) long night conditions: N = 60 (control), 66 (*Nca*^KO1^), interaction between stimulus and genotype: p = 0.0472. Performance does not differ between the genotypes in (H) short night conditions: N = 61 (control), 61 (*Nca*^KO1^), interaction between stimulus and genotype: p = 0.2395. Two-way ANOVA with Dunn’s multiple comparison tests. Error bars: SEM.

Taking advantage of this finding, we quantified mechanosensory habituation in control and *Neurocalcin* knockout flies entrained to short or long night conditions (Fig. 3F). Strikingly, while mechanosensory habituation was reduced in *Neurocalcin* knockout flies compared to controls under long night conditions (Fig. 3F), this difference was abolished under short night conditions (Fig. 3G). These data strongly suggest that Neurocalcin regulates habituation directly through its effect on night sleep.

### Loss of Neurocalcin decorrelates night length and synaptic protein expression

Intriguingly, within-genotype comparisons revealed that control flies exposed to short nights exhibited a reduced rate of mechanosensory habituation compared to control flies exposed to long nights (Figure S4A), indicating that limiting night sleep (Fig. 3C) disrupts mechanosensory habituation during the following day. Conversely, *Neurocalcin* knockout flies, which exhibit improved night sleep in short versus long nights (Fig. 3C), exhibited the opposite correlation: an enhanced rate of habituation in short versus long night conditions (Figure S4B).

How might night duration modulate mechanosensory habituation occurring after nighttime sleep? Prior studies have shown that sleep deprivation (SD) in adult flies impairs synaptic downscaling, resulting in increased levels of the presynaptic active zone protein Bruchpilot ^43^ that can be visualised through immuno-fluorescent microscopy. Prolonged (16 h) SD was initially shown to increase presynaptic Bruchpilot expression by ∼ 75% across *Drosophila* brain neuropil regions ^44^. More recently, 12 h night-specific SD was shown to modulate Bruchpilot to a more limited degree, enhancing Bruchpilot expression in cholinergic neurons by ∼ 25%, and in GABAergic neurons by ∼ 10% ^45^. Loss of the *Drosophila* Insomniac protein, which reduces overall day and night sleep by ∼ 65% in male flies ^32^, also leads to a 40-50% increase in presynaptic Bruchpilot expression ^46^.

Extending our analysis in the context of the synaptic homeostasis hypothesis ^5^ and the above studies, we investigated whether altering night length was sufficient to modify presynaptic Bruchpilot expression in control flies, and if so, whether loss of Neurocalcin affected this relationship. We immuno-labelled synaptic Bruchpilot in isolated brains of control and *Neurocalcin* knockout flies following three days of entrainment to short or long night conditions, then quantified fluorescent intensity of synaptic Bruchpilot through confocal microscopy, normalising fluorescence to a control DNA marker (DAPI, see Methods).

Consistent with the relative subtlety of this manipulation compared to mechanical sleep deprivation ^44,45^ or loss of Insomniac ^46^, we observed a trend towards a small increase (∼ 16%) in Bruchpilot throughout the brain neuropil region of control flies subjected to short versus long nights, which was non-significant (Figure S4C). However, in the neuropil-rich antennal lobe regions, which process olfactory information in insects, we observed a significant increase (∼ 40%) in synaptic Bruchpilot levels in short versus long night conditions (Fig. 4A,B).. Notably, we did not observe a similar increase in antennal lobe Bruchpilot expression in *Neurocalcin* mutants exposed to short versus long nights (Fig. 4C,D), nor in whole-brain Bruchpilot immuno-fluorescence (Figure S4D), suggesting that the impaired sleep quality of *Neurocalcin* exposed to long nights (Fig. 3A,C) blocks enhanced synaptic downscaling that occurs in wild types flies under this condition.

**Fig. 4.**
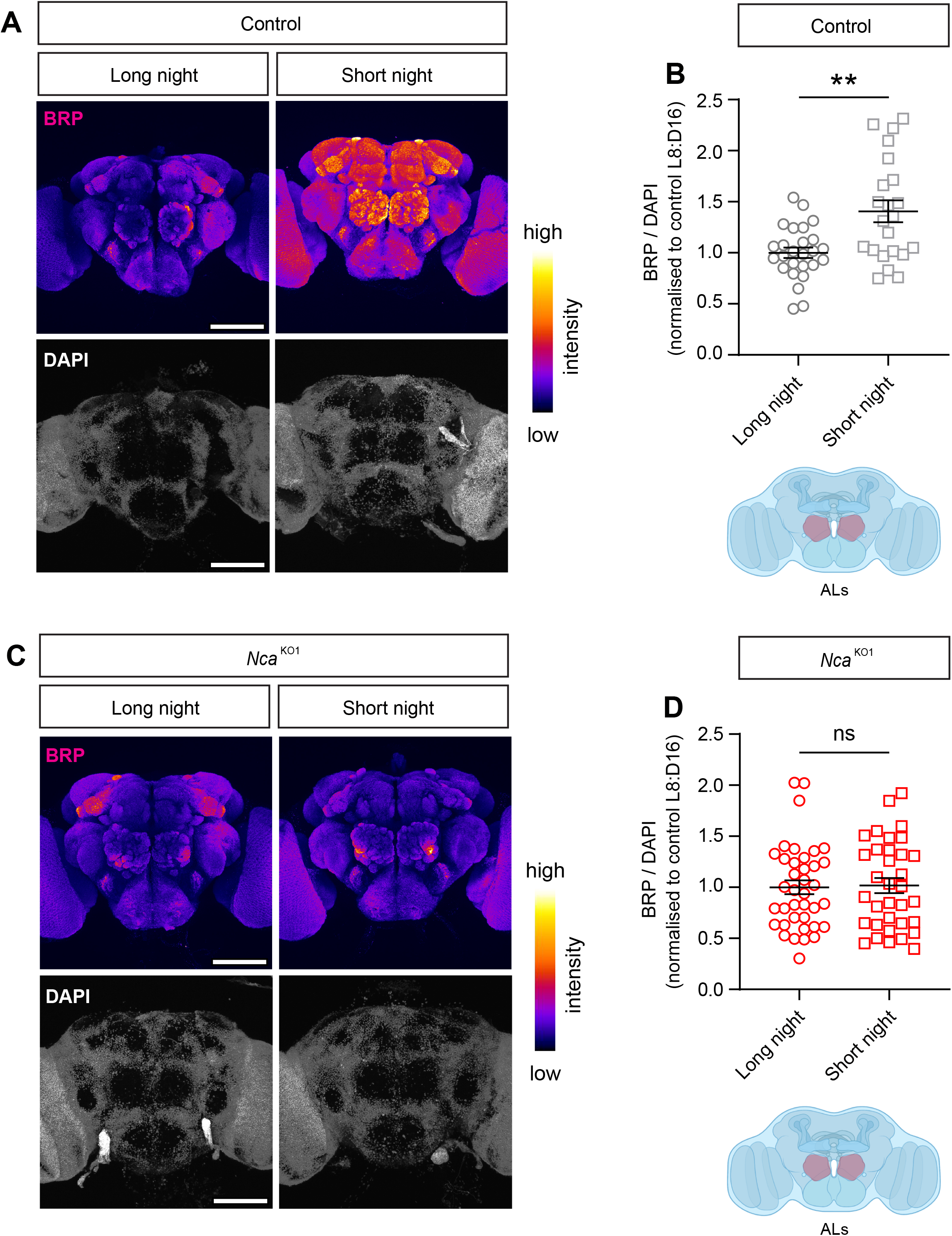
Night sleep duration modulates Bruchpilot immunofluorescence in sensory circuits. (**A, C**) Representative images of (A) iso31 and (C) *Nca*^KO1^ whole brains immuno-stained for anti-BRP (top), with a DAPI counterstain (bottom), in long night (L8: D16) vs short night (L16: D8) conditions. Scale bars: 100μM. (**B, D**). Mean BRP immunofluorescence within antennal lobe neuropil regions (two data points per brain), normalised to whole-brain DAPI immunofluorescence, is increased in short-versus long-night conditions in controls (p = 0.0018, unpaired t-test with Welch’s correction), but unchanged in *Nca*^KO1^ brains (p = 0.81, Mann-Whitney U-test). Values are normalised to the mean value in long night conditions. Error bars: SEM.

Collectively, the above data support a model in which reduced night sleep in control flies exposed to short nights perturbs synaptic downscaling in specific neuropil domains compared to flies experiencing long nights. We hypothesise that such domains may include circuits relevant to mechano-sensation and movement, where impaired downscaling could limit the ability of habituation-promoting mechanisms (such as enhanced synaptic transmission from local inhibitory neurons ^23^) to dampen functional connectivity between mechanosensory neurons and downstream motoneuron outputs. The improvement of sleep quality in *Neurocalcin* mutants exposed to short nights (Fig. 3A,C-E) breaks the normal correlation between night length and synaptic Bruchpilot expression (Fig. 4), leading to improved rather than impaired mechanosensory habituation in short compared to long night conditions (Figure S4).

Further studies will be required to test this model – in particular, by identifying mechanosensory and/or pre-motor circuits in which synaptic output is modulated by habituation-promoting mechanisms. Intriguingly, recent work has shown that continuous or repetitive mechanosensory stimulation, akin to the gentle rocking of a human baby, increases daytime sleep in *Drosophila* through a habituation-like mechanism ^47^. In concert with our findings, these data suggest an intertwined and potentially bi-directional relationship between mechano-sensation, sleep, and habituation, across both day-night and seasonal cycles. Finally, we note that sleep and habituation are dysregulated in a variety of human neurodevelopmental disorders ^48-52^ and corresponding animal models ^49,53-57^. Since habituation promotes attention to environmental novelties that form the substrates of memory-related coincidence detection mechanisms ^58^, our work has implications for understanding the common co-morbidity of sleep, habituation, and memory defects in these clinically important conditions.

## ACKNOWLEDGEMENTS

We thank Dr. Gabriel Aughey for comments on the manuscript, members of BKFLab for technical assistance with the DART system, and members of the Zantiks team for technical assistance with the Zantiks MWP system. This study was funded by a MRC Senior Non-Clinical Fellowship (MR/V03118X/1) and a BBSRC Research Grant (BB/R02281X/1) to J.E.C.J; a BBSRC Research Grant (BB/X00094X/1) to S.L and J.E.C.J; a Wellcome Early Career Fellowship to A.D.W (308298/Z/23/Z); and a BBSRC Research Grant (BB/W014939/1), and a Startup and CRE fund from the Division of Genetics and Genome Biology of University of Leicester (RM33077K), to K-F.C.

## AUTHOR CONTRIBUTIONS

S.A.L – Conceptualization, Methodology, Investigation, Writing - Original Draft, Writing – Review & Editing, Visualization, Funding acquisition; A.D.W – Methodology, Investigation, Writing - Original Draft, Writing – Review & Editing, Visualization. K-F.C – Investigation, Writing – Review & Editing. J.E.C.J – Conceptualization, Investigation, Visualization, Writing - Original Draft, Writing – Review & Editing, Supervision, Project administration, Funding acquisition.

## MATERIALS AND METHODS

### Experimental model and subject details

Experimental models were fruit flies of the species *Drosophila melanogaster*. Genotypes used were iso31 (an isogenized strain of *w*^1118^ flies and a kind gift from Prof. Kyunghee Koh, Thomas Jefferson University), *w*^1118^;+; *Nca*^KO1^/TM2, and *w*^1118^;+; *Nca*^KO2^/TM2 ^9^ To control for genetic background, these lines were outcrossed into the iso31 background for 5 generations. Flies were raised and maintained on standard fly food under 12 h: 12 h light-dark cycles (12L: 12D) at a constant 25°C. Experiments took place at 25°C, and unless otherwise indicated, in 12L: 12D conditions. Experiments were conducted on male flies.

### Habituation protocol

Drosophila ARousal Tracking (DART) analysis was performed as described previously ^1^. Briefly, five-to seven-day old male flies were individually loaded into glass behavioural tubes (Trikinetics inc., MA, USA) containing 4% sucrose and 2% agar, and left for 24 h to acclimatise in the indicated L:D conditions. The mechanical stimulus consisted of a mechanical vibration in the form of a train of five 200 ms pulses separated by 800 ms intervals, set at the DART system’s maximum intensity, which elicited a robust movement response. The habituation protocol consisted of 20 mechanical stimuli delivered at ten-minute intervals, beginning at Zeitgeber time 3 (ZT3). Videos were taken using a USB webcam (Logitech). Speed (mm / s) was analysed by the DART system from absolute position, and binned in one min intervals. Flies were excluded from analysis if they exceeded an average of 1 mm / s speed in the five one-minute bins preceding the first stimulus. Response to the stimulus was defined as the mean speed (mm / s) in the one-minute bin following the stimulus. Speed was normalised within genotype to the mean of the response to the first stimulus. To derive the % of flies responding to stimuli, a “response” was distinguished from background movement by the (arbitrary) threshold of a change in mean speed between the one-minute bins preceding and following a stimulus of ≥10% of the mean response to the first stimulus.

A distinct automated *Drosophila* tracking system, the Zantiks MWP unit (Zantiks Ltd., Cambridge, UK) was used to generate an equivalent habituation protocol. 5-7 day old male iso31 control flies were loaded into individual wells filled with 4% sucrose and 2% agar in a 48-well plate, and left to acclimatise for 24 h in a 12 h light: 12 h dark cycle at 25°C. *Drosophila* were loaded into the Zantiks unit at ZT0, and the habituation protocol commenced at ZT3. The habituation protocol again consisted of 20 mechanical stimuli delivered at 10 min intervals, each stimulus consisting of a single 1 s pulse set at medium strength, generated by the Zantiks mechanical stimulation stand. Distance travelled is automatically tracked by the Zantiks system, and total distance travelled was binned in 1 min intervals. Scripts used for the habituation protocol on the Zantiks MWP are available at github.com/abigaildwilson/zanscripts.

### Sleep analysis

Sleep analysis was performed using the *Drosophila* Activity Monitor 5 system (DAM5H, Trikinetics inc., MA, USA), which improves on the fidelity of the earlier DAM2 system by using 15 infrared beams per tube, rather than a single one, to track movement ^2^. As above, three-to five-day old male flies were loaded in behavioural tubes, and were left for two full days to acclimatise at the relevant light cycle. On the third day the number of beam breaks was measured using the *Drosophila* Activity Monitor. Sleep was defined using the standard definition of > 5 mins of inactivity, which has been shown to correlate with postural changes, increased arousal threshold and altered brain activity patterns ^2^. Sleep bout number and duration was analysed using the open-source software suite phaseR (https://github.com/abhilashlakshman/phaseR/tree/main) ^3^.

### Immuno-histochemistry and confocal microscopy

Brains were dissected in phosphate-buffered saline (PBS) (Sigma Aldrich) between ZT1-2, fixed by 20 min room temperature incubation in 4% paraformaldehyde (MP biomedicals) diluted in PBS + 0.3% Triton-X (Sigma-Aldrich) (PBT), washed three times in PBT, and blocked for 1 h in 5% normal goat serum in PBT. Brains were incubated for 48 hours at 4°C in primary antibody (mouse anti-BRP [nc82], Developmental Studies Hybridoma Bank, at a concentration of 1:200); then washed three times in PBT and incubated overnight at 4°C in secondary antibody (goat anti-mouse AlexaFluor 647, ThermoFisher Scientific, at a concentration of 1:500) and counterstain (DAPI, ThermoFisher Scientific, at a concentration of 1:500). Brains were washed three final times before mounting in SlowFade Gold anti-fade mountant (ThermoFisher). Images were taken with a Zeiss LSM 980 confocal microscope with an EC ‘Plan-Neofluar’ 20x/0.50 M27 air objective, taking z-stacks through the entire brain with step sizes of 1 μm. Images were visualised and analysed using ImageJ; maximum intensity z-stack projections were generated, and following background subtraction, regions of interest (ROIs) were manually drawn and mean pixel intensity calculated. BRP intensity readings from both the whole central brain and limited to the antennal lobes were normalised to DAPI readings from the whole brain. Data is pooled from three separate experiments, with brains within experiments dissected, immuno-stained and imaged in parallel. Values are normalised to the mean values of the long night (L8: D16) condition within an experiment.

## SUPPLEMENTARY FIGURE LEGENDS

**Supp. Fig. 1**. (**A**) Quantification of mean locomotor speed over time in response to a single mechanical stimulus. Mean speed of N = 11 control (iso31) male flies are shown. (**B-D**) Mean speed over time as a further single mechanosensory stimulus is delivered following an interval of (B) 1 h (N = 87), (C) 4 h (N = 36), or (D) 24 h (N = 27). (**E**) Mean speed in the 1 min bin following a single stimulus at the indicated interval after the initial habituation protocol. Data are normalised to the naïve response to the first stimulus in the 20-stimulus train (N = 87). Responses progressively return towards the baseline level. (**F**) Schematic of the Zantiks system. Individual flies are loaded into a 48-well plate containing a solidified agar and sucrose food surface. Mechanical stimuli are delivered via a motor beneath the plate, with locomotor responses recorded via videography. (**G**) Mean locomotor responses over time of N = 24 flies to a single mechanical stimulus (solid line), quantified as distance travelled in a 1 min interval. (**H-I**) Mean distance travelled per 1 min (**H**; line represents mean value over time), and mean distance travelled in the 1 min following each mechanical stimulus (**I**), in N = 48 control (iso31) male flies exposed to 20 successive mechanical stimuli in the Zantiks system. Error bars in E, I: SEM

**Supp. Fig. 2**. (**A**) Mean speed of locomotion over time, and (**B**) mean speed in one-minute bin following each stimulus of control (iso31) (N = 41) and a second independently generated *Nca* knockout line (*Nca*^KO2^) (N = 30) adult male flies exposed to 20 consecutive mechanosensory stimuli. Data are normalised within each genotype to the naïve response. *Nca*^KO2^ flies habituate more slowly than controls (interaction between stimulus number and genotype: p < 0.0001, two-way ANOVA). Error bars: SEM. (**C**) Mean speed of control and *Nca*^KO1^ adult male flies in the one-minute bin following a single mechanosensory stimulus of varying strength (values are normalised to the strength used in the habituation assay). Control: N = 22-52; *Nca*^KO1^: N = 15-41. *p < 0.05, ns: p > 0.05, Mann-Whitney U-test.

**Supp. Fig. 3**. (**A**) Schematic illustrating components of the *Drosophila* Activity Monitor 5 (DAM5) system. Flies are individually housed within glass behavioural tubes. Locomotion is tracked via breaking of 15 infrared beams across the tubes. Sleep is derived from periods without locomotion (see Methods). (**B-D**) Total minutes of sleep during subjective day (B), number of individual sleep bouts during subjective day normalised to hours of day (C), and mean length of sleep bouts occurring during the day (D), for populations presented in Fig. 3A-B. Kruskal Wallis test with Dunn’s multiple comparison test. *p < 0.05, **p < 0.01, ***p < 0.001. Error bars: SEM.

**Supp. Fig. 4**. (**A-B**) Within-genotype comparisons of the effect of light cycle on habituation. (A) Iso31 flies habituate more quickly in long-night conditions (interaction between genotype and light cycle: p = 0.0027). In contrast, *Nca*^KO1^ flies (B) habituate more quickly in short-night conditions (interaction between genotype and light cycle: p = 0.0494). (**C-D**) Mean Bruchpilot (BRP) immunofluorescence within the whole central brain, normalised to DAPI, is not significantly changed between short-versus long-night conditions in control (C: p = 0.1648), or *Nca*^KO1^ brains (D: p = 0.5461). Unpaired t-test with Welch’s correction (C) or Mann Whitney U-test (D). Values are normalised to the mean value in long night conditions. Error bars: SEM.

