## Supplementary figures and images for "Nighttime sleep promotes mechanosensory habituation in *Drosophila*"

### FIGURE S1

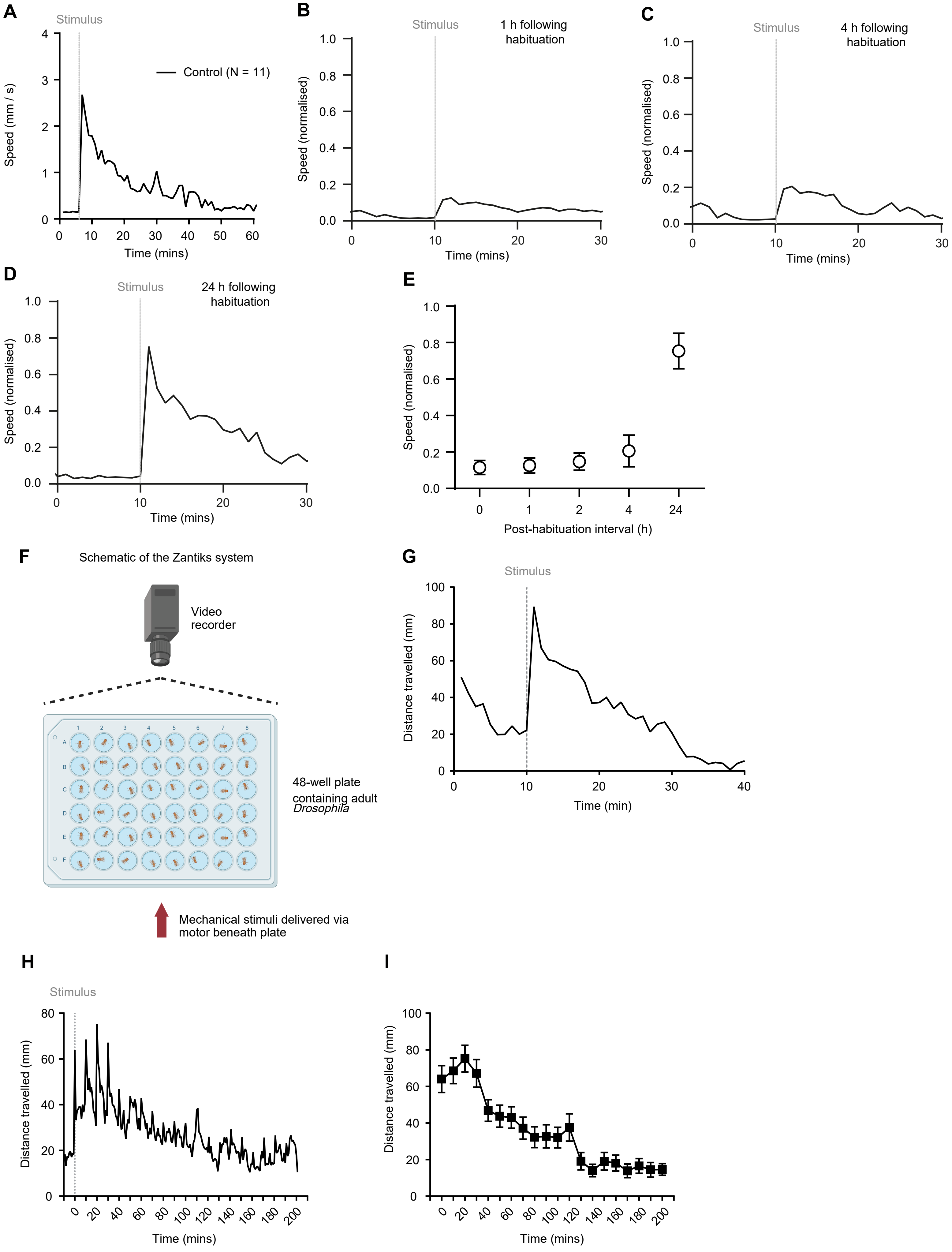

### FIGURE S2

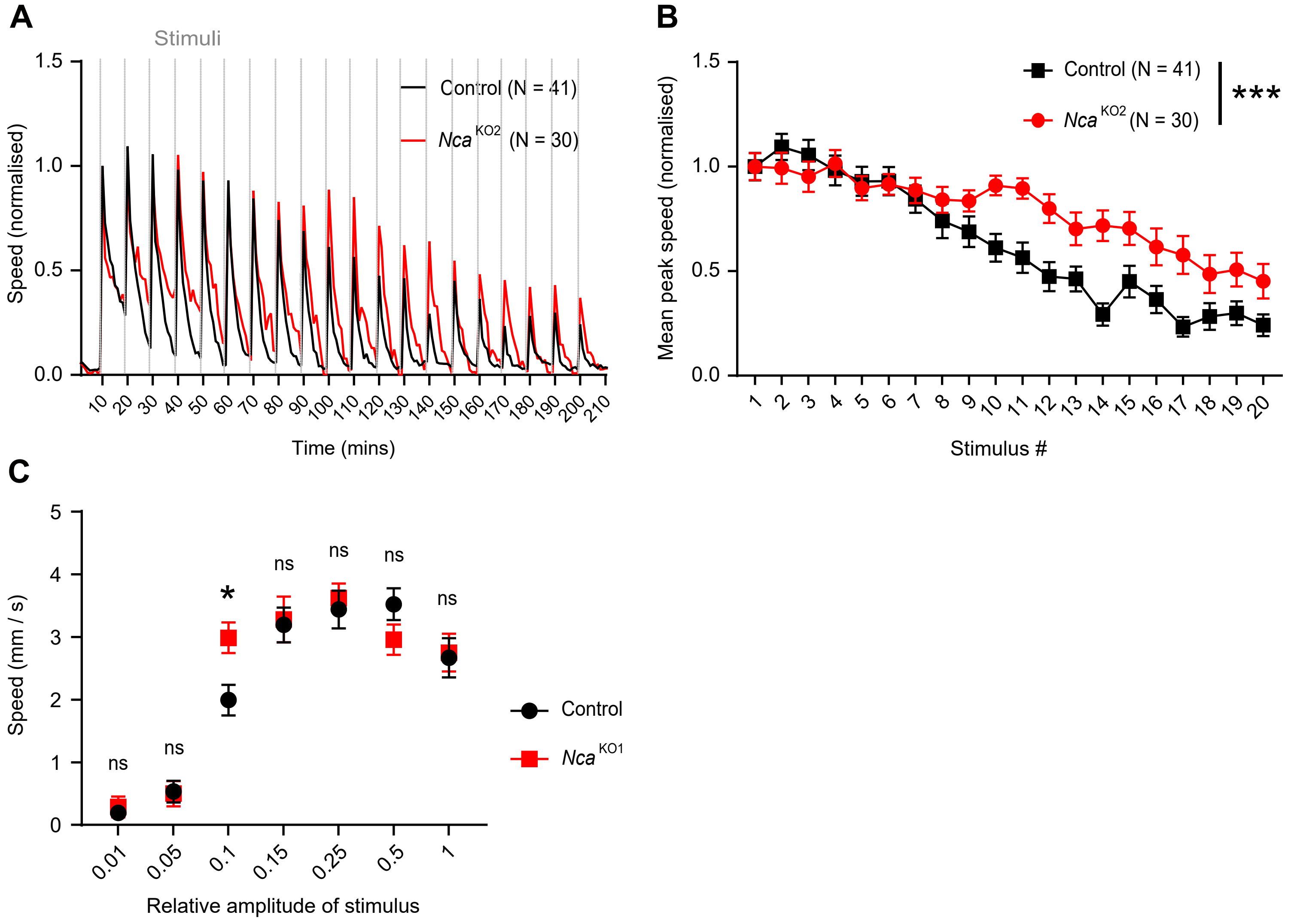

### FIGURE S3

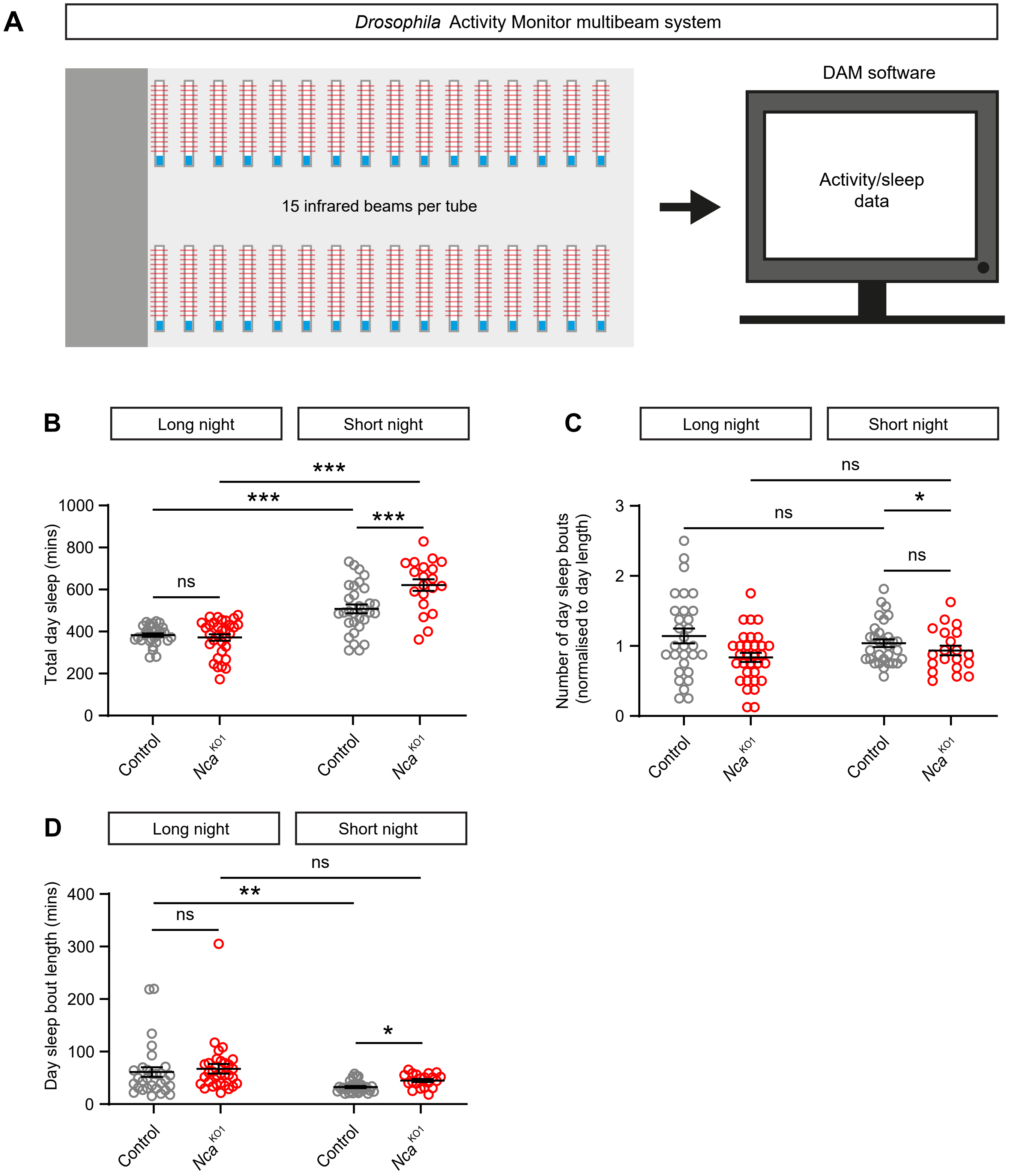

### FIGURE S4

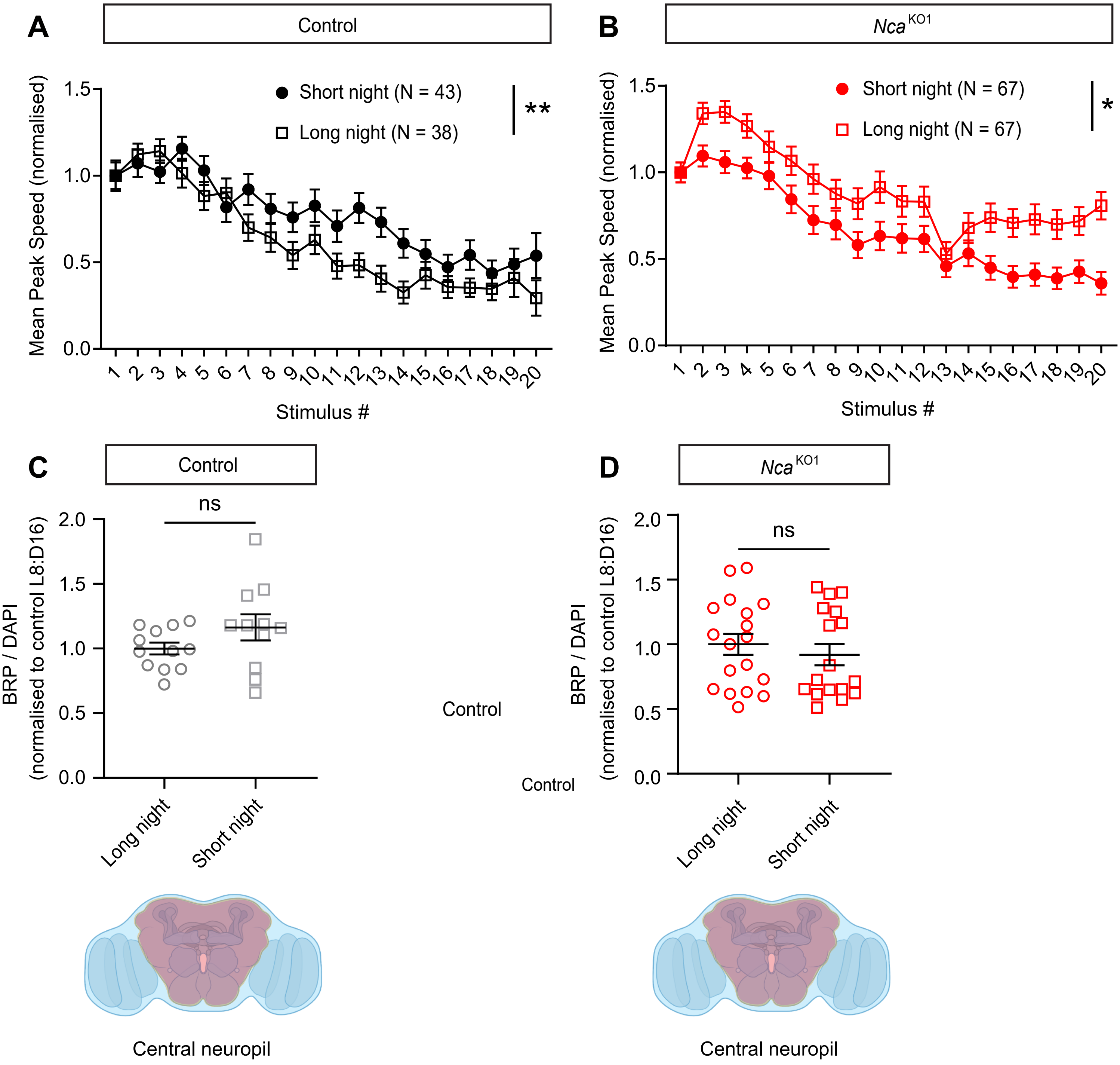
